# KmerFinderJS: A client-server method for fast species typing of bacteria over slow Internet connections

**DOI:** 10.1101/145284

**Authors:** Jose Luis Bellod Cineros, Ole Lund

## Abstract

**Motivation:** KmerFinder is a program based on K-mer statistics for identifying bacterial species in whole genome data, that as a web server that have been used more than 10.000 times. KmerFinderJS is a development of the KmerFinder that benefits from the downsampling of data using a prefix filtering used by KmerFinder, to minimize amount of data that needs to be transferred between the client and the server.

**Results:** KmerFinderJS replaces the python based hash structure for holding the databases with a true Key-value database. These improvements are shown to lead to a many-fold speed up of species identification with the internet transfer speeds that are realistic to expect today. It is also shown that the method can find the true content of an artificial metagenomic cocktail with no false positives.

**Availability:** The method is freely available at https://cge.cbs.dtu.dk/services/KmerFinderJS/ and as a source code at https://bitbucket.org/genomicepidemiology/kmerfinderjs

**Supplementary information:** Supplementary data are available at *biorxiv* online.

## 1 Introduction

KmerFinder (Larsen, Mette V., et al. (2014)) is a method developed for prediction of bacterial species. The method is based on the sharing of a high number of sequence fragments between an input query file with an unidentified species and a reference database. The fragments result from the split of the sequence into substrings of fixed length (K-mer of length 16). The sequence with the highest numbers of K-mers shared with the sequence query will be the predicted species. To reduce database size and search time only K-mers that start with a given prefix are used in the comparisons. The standard prefix used for bacterial identification is the 5mer “ATGAC”. A so called winner-takes-all option (Hasman, H, et al. (2013)), is implemented that iteratively finds the hits by in each round only keeping the top hit and removing all K-mers found in that hit from the dataset before the next search.

A major bottleneck for the speed of KmerFinder is that the input data is typically 200mb, and the databases in the order of 1Gb. Transfer of the smaller of the databases over a typical internet connection will take ~12 minutes (with a speed of 1Gb/h connection) which then becomes a lower bound for the runtime of the program. This is an example of the I/O bottleneck, which is one of the biggest problems in the Big Data ecosystem where huge databases need to get closer to the computing resources. Several new methods like, faster network connections, data-compressing algorithms, parallel computing, new data representations and data streaming have been considered for solving this problem (Stephens, Z. et al. (2015)) In the work of (Gautier, L. and Lund, O. (2013)) only K-mers from a random selection of reads were sent to the server which had a very large database containing gen-bank and all available completed genomes. Another approach was pursued by (Saputra, D. et al. (2015)) which constructed a small database (~5mb) of informative 50mers form 16s sequences, which could be send to the client where it was matched against the input data. Here we present a new implementation of KmerFinder. Several key features have been developed: a complete new JavaScript implementation, client-server architecture and centralized database. We take advantage of that the 5mer prefix downsamples the data 4^5^=1024 times, and if this downsampling is done on the client side it drastically reduces the amount of data that needs to be send to the server, and thereby the transfer time to seconds.

## 2 Methods

KmerFinder (Larsen, M. et al. (2014)) is a species prediction algorithm based on the number of shared K-mers (16-mers) between a sequence file and a reference database. It was first implemented as a Python program and later expanded with a scoring-scheme called winner-takes-all (Hasman, H, et al. (2013)). In this new implementation, the scoring algorithm is split into two parts, client and server and executed in a web environment accessible via a web browser.

The algorithm consists on the following steps: first the extraction of the overlapping 16-K-mers from the sequence file, second the comparison of the query set of K-mers with the reference database of sequences and third, the process of finding matches (i.e. candidate templates for the input file) from the reduced database sent to the client. KmerFinderJS starts the execution on the client; the user accesses the file-browsing interface of his operative system either by drag and drop on the web-interface or by clicking on it. Once the sequence file are added, the program will immediately start extracting overlapping 16-mers and storing them in a JavaScript object with the key being the 16-mer and the value the number of times the K-mer was seen in the file.

After parsing the file, the object is sent to the server using a GET HTTP request. The server receives the object and extracts all keys, querying the database for each K-mer and finds all templates that belong to that query. This will create a reduced version of the database that contains only the K-mers present in the query and its associated templates.

The reduced database is sent back to the browser where it is sorted in decreasing order based on the number of K-mers that each template has. The first template according to this sorting scheme will be the candidate that the algorithm chooses as the possible species for the input file. Next the K-mers associated to this template will be removed from the input object to speed up the search (this template will not appear in further iterations since the input object doesn’t share any K-mers with it). The reduced database containing the templates from the first round is saved for later comparison. This continues in an iterative fashion where all K-mers from the input query will be compared with the K-mers in the initial reduced database creating a new template database which is sorted in decreasing order based on the number of shared K-mers. The first template is then considered the second candidate for a match. This process will be repeated until 100 templates have been found or the next template found has a p-value above 0.05. The above algorithm for finding the next best hits is called the “Winner-takes-all” approach. (See Supp. materials Figures 1 and 2)

## 3 System Design

The new implementation is entirely made in JavaScript using the EcmaScript 6 standard (ES6) in a NodeJS environment. The code was tested first in the server without relying on any web browser. Only recent versions of major Browsers support ES6 (http://www.ecma-international.org/ecma-262/6.0/) so a two-step transpiling process was used, first to produce ES5 code from ES6 using a tool called Babel (https://babeljs.io) and then converting the code developed on the server to a compatible browser-version using Browserify (http://browserify.org) The python implementation of KmerFinder uses a python dictionary stored in a pickle file to load the reference database each time the program is executed, increasing the running time and has the potential problem of overloading the main memory of the server. KmerFinderJS uses Redis (https://redis.io), a centralised in-memory database, to store the reference database. Redis stores data in key-value pairs. Keys are identified as string, and values can be Lists, Sets or Hashes of Strings, among others. For our database of sequencing files we store the unique K-mers as keys and the value for each of K-mer is a list of strings that encode individual templates.

## 4 Results

To benchmark KmerFinderJS with the implementation in python, a dataset of 207 reads from the NCBI archive was used to time the total time of execution, the time to extract the overlapping K-mers and the time to query the database. This is the same dataset used in (Thomsen, M. et al. (2016)) to benchmark the MLST method. KmerFinderJS was ran as command-line program to compare with the running time of the KmerFinder in python script, both running on a MacBook Pro 2.2 GHz Intel Core i7, and the time of sending data over the network was added to the total time based on different speed connections. The average time for all files was calculated to compare the two methods. The main bottleneck of the method is the extraction of the K-mers due to the fact that we need to parse the whole file sequentially. Both KmerFinderJS and KmerFinder showed similar running times, with KmerFinderJS being faster overall except on the winner-takes-all step. It should also be noted that Redis ran in a Docker container so a native installation could perform better. The impact of avoiding sending the whole file to the server clearly lowers the running time of the method. KmerFinder needs to send the whole file over the network, which for the fastest Internet connection is around 20% slower than KmerFinderJS, taking around 1 minute. (See Supp. materials S1 figures 3 to 6). A real test running KmerFinderJS on the browser was performed in Moshi, Tanzania, at the Kilimanjaro Christian Medical College using a 3G mobile phone connection. For this test a metagenomic sample was used (Peabody, M. (2015)). The first step of extracting the overlapping K-mers took 26.4 seconds. Sending the K-mers and receiving the reduced database took 1.6 minutes and finding matches on the reduced database took 16.9 seconds KmerFinderJS clearly benefits from only sending the overlapping K-mers and the reduced database and the total time remains almost constant for all speed connections. (See Supp. materials figures 7 and 8).

